# Linking neural representations to behavior using generalization

**DOI:** 10.64898/2026.04.21.719991

**Authors:** Miguel Angel Nuñez-Ochoa, Fengtong Du, Lin Zhong, Scott Baptista, Michalis Michaelos, Alex Sohn, Liad Baruchin, Sylvia Schröder, Carsen Stringer, Marius Pachitariu

## Abstract

Sensory-guided decisions are the result of sensorimotor transformations across many brain areas. Recent studies have localized the motor- and decision-related components of these transformations using brain-wide neural recordings. It has been more difficult to localize sensory computations in the same way. Here we developed a new approach for linking sensory computations to behavior by training mice to discriminate between two stimuli and testing their responses with new stimuli. In separate animals, we calculated the similarity of neural representations between train and test stimuli, using recordings of up to 73,000 simultaneously-recorded neurons from 9 primary and higher-order visual areas (HVAs) across layers 2 and 3. We found that neural discrimination on test but not train images correlated with behavioral discrimination, and this relation required prior visual experience as it was not present in dark-reared mice. The link between neural and behavioral performance was highest in the medial HVAs, suggesting this region is a critical component of sensory transformations and generalization.

## Introduction

One of the main goals of systems neuroscience is to understand the neural computations that give rise to animal behaviors. This requires multiple steps, such as identifying the brain regions involved, determining the transformations they perform on their inputs, and determining how the resulting abstract representations guide behavior [1–14]. In this process, sensory transformations play a critical role in generating actionable, high-level visual representations from low-level sensory features [15–17]. Unfortunately, it is often difficult to identify where such sensory processing takes place in the brain, due to the additional non-sensory processing that intervenes between a sensory computation and the resulting behavior [18–20]. These non-sensory computations have an influence, not just on behavior, inducing trial-to-trial variability, but also on neural activity across the entire brain [21–24]. Due to the pervasive influence of internal states and motor signals on neural activity, it is typically difficult to identify the direct effects that sensory computations have on behavior.

A common approach to linking sensory processing to behavior has been to parametrically modify the difficulty of a task by varying the intensity of a stimulus. This produces psychometric discrimination curves, which can be compared to neurometric discrimination curves obtained as a measure of neural discrimination from a specific neuron or brain region [25, 26]. If the two have similar shapes and slopes, one may tentatively infer a relation between the respective brain area and behavior [27]. However, this approach has encountered conceptual obstacles, as the neural information available in sensory brain regions has been shown to far surpass the information that is available for behavior [28]. More generally, stimuli used for discrimination in animal tasks are often highly simplified and only cover a small region of the neuro-behavioral space.

Here we introduce a different approach for identifying sensory transformations that lead to behavior. Rather than training animals closer to psychometric thresholds, which may be impossible, we distinguish between a “training” phase with one set of stimuli, and a “testing” phase with a different set. This is similar to machine intelligence training tasks, where trained models are always evaluated on out-of-sample test data, to determine the true performance of that model. We created an ensemble of train/test stimulus combinations that produced different *levels* of generalization [29–31] by natural variation in the higher-order similarities between the train/test images. We then took advantage of the different levels of behavioral generalization of mice across these stimuli to compare behavioral responses to neural responses and identify brain regions where these match.

## Results

To illustrate the relation between sensory neural computations and behavioral discrimination, we describe a simplified toy model of sensory-based decision making (Figure 1a). We then use this model to predict what aspects of behavior *may* correlate with neural activity, and design experiments that test these predictions. In the model, two training stimuli drawn from different distributions are represented as points in neural population space (Figure 1b). The model learns a decision boundary that bisects these two points. Since two points can always be separated, the classification accuracy on training data is always 100% regardless of what distributions the points were drawn from. This situation mimics neuroscience tasks in which animal behavior is trained and evaluated on the same stimuli. While animals do not necessarily perform at 100% accuracy in such tasks, that is likely due to motor and behavioral state variability, rather than variability in sensory neural computations. Thus, when neural and behavioral discrimination are compared to each other, there is little to no relation present (Figure 1c and see [28]).

**Figure 1:**
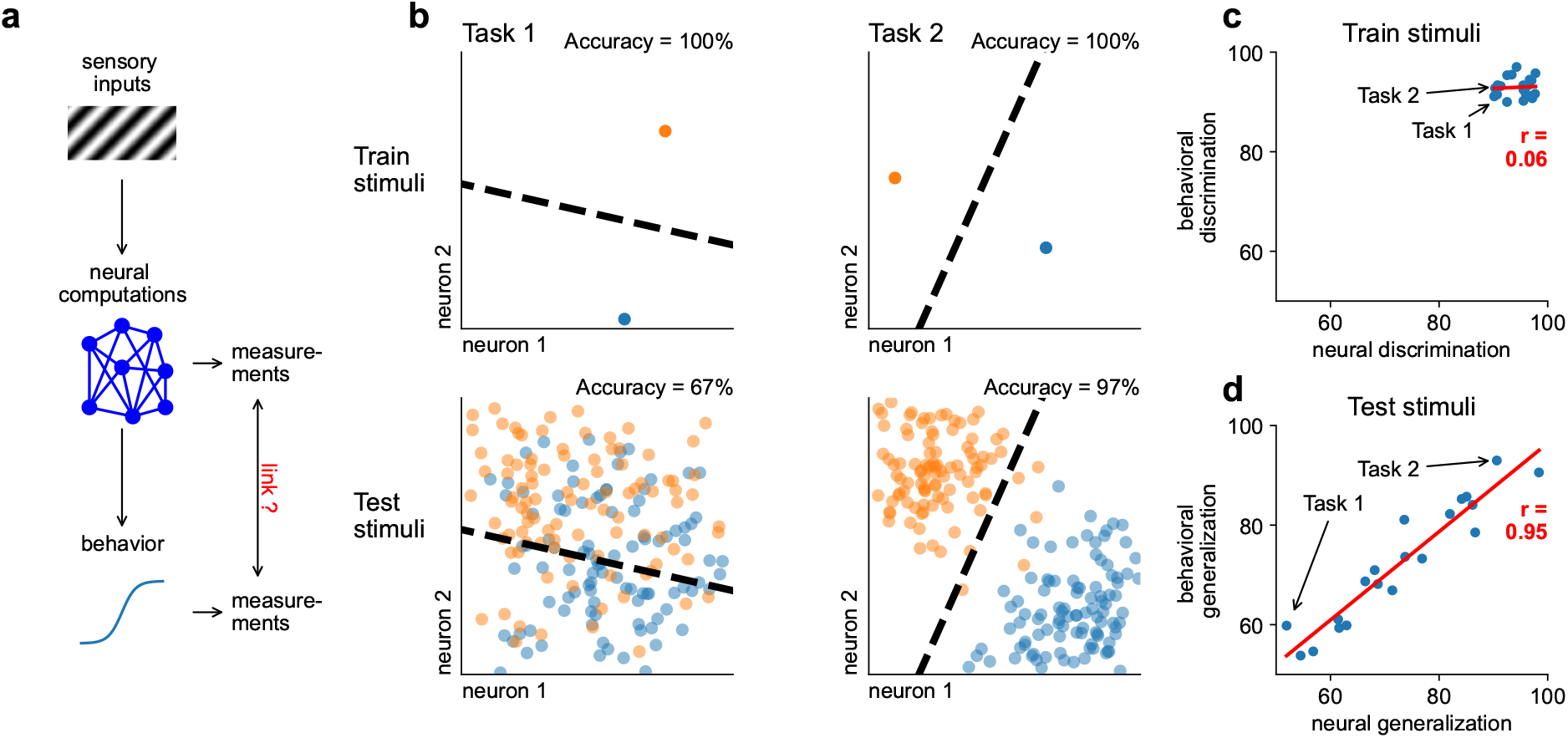
Predicted relation between behavioral and neural discrimination. **a**, Schematic of sensory processing, decision-making and experimental measurements. **b**, Simulated distributions for two stimulus categories in two different tasks (left and right columns). Top: Single stimuli from these categories used to learn a discrimination boundary (dotted line) on train trials. Bottom Test samples from the distributions overlaid with the discrimination boundary from top row. **c**, Schematic relation between neural and behavioral discrimination on train trials, assuming perfect performance (illustrated in **b** top row) with added motor and neural noise. **d**, Same as **c** on test trials.

To avoid this limitation, we may instead study discrimination performance on “test” stimuli, not used for training (Figure 1b). In this case, the discrimination accuracy can be less than 100% and possibly down to chance depending on the neural distributions. Critically, the accuracy more closely aligns with the true separation between the distributions of neural responses. Thus, discrimination performance on test stimuli results in a correlation between neural and behavioral generalization (Figure 1d). Below, we illustrate how this approach can be implemented to study animal behavior and its relation to neural activity.

### Behavioral generalization

First, we estimated behavioral discrimination accuracy on test stimuli. To obtain a sufficient number of sessions with different training and testing images, we designed a visual task that could be learned quickly (Figure 2a). Headfixed mice, running on a treadmill, had to discriminate between two stimuli that spanned a large field of view (270 degrees). Stimulus contrast was increased and decreased linearly as a function of the running speed of the mice. Thus, simply by running forward, mice initiated the trials themselves, which is advantageous for fast training. Images were alternated in pseudo-random fashion, for a total of 400 trials per day. Each mouse was trained for five days on a pair of images, with the fifth day including test images from the same categories A and B. We used stimuli from eight categories (leaves, circles, dryland, rocks, tiles, squares, round leaves, and pavement) (Figure 2a). Most pairs of stimuli were shown twice, in different mice (Figure 2b). Mice were trained on a new pair of images each week, with some mice completing 9 weeks of training (Figure 2c, Figure S1).

**Figure 2:**
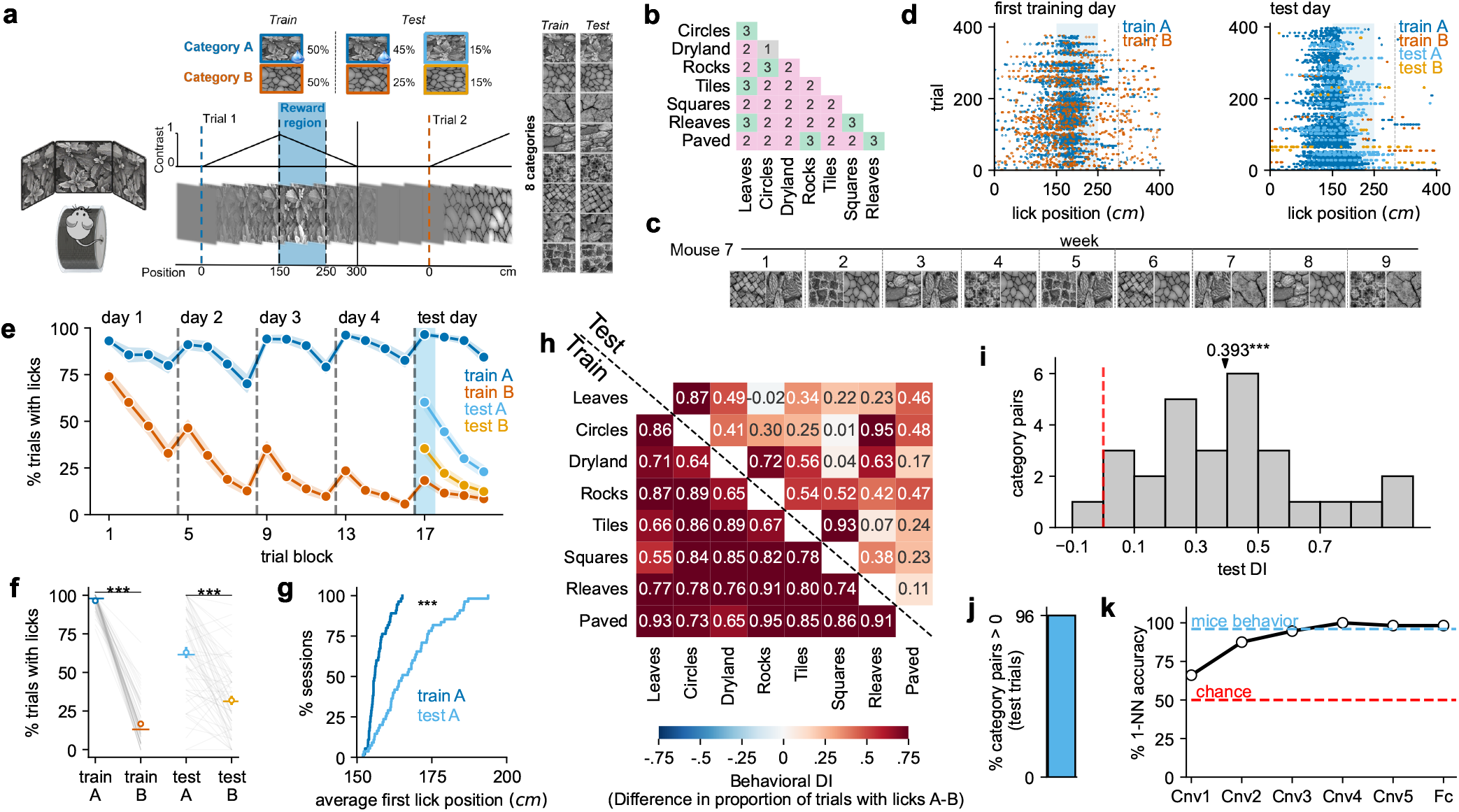
Behavioral performance on train and test stimuli. **a**, Task schematic denoting an abstract position variable linked to the running speed of the mouse that is used to advance through increasing and decreasing contrast of natural images displayed in a 270 degree field of view. (right) All images used across experiments. **b**, The number of repetitions of each pair of train images in different mice. **c**, Training stimulus pairs for one mouse. **d**, Lick rasters from an example mouse on the first training day (left) and on the test day (right). **e**, Fraction of trials with lick behavioral responses. Each day is split into 4 equal blocks of trials for visualization, the first block (shaded) is the block used to analyze the test session (n = 52 sessions in 11 mice). Error bars represent s.e.m. across sessions. **f**, Fraction of trials with lick behavioral responses per pair of trained and test images (n = 62 sessions in 15 mice), paired t-test). **g**, Cumulative distributions of first lick position in the reward region for train and test A trials (n = 62 sessions in 15 mice, Cramér-von Mises test). **h**, Difference of lick rates between A and B trials, denoted “discrimination index” (DI), for both train and test images, where the test DI is normalized by the train DI (see Methods). **i**, Distribution of test DI from **h** (n = 62 sessions in 15 mice, one-sample t-test). **j**, Fraction of pairs with positive test DI from **i. k**, Comparison of mouse generalization behavior on test images to nearest-neighbor classification using different layers of Alexnet.

Each week, mice started with high response rates to both training stimuli, and gradually reduced their licking to the non-reward stimulus, both within-day and across days, with clear signs of learning emerging from day 1 (Figure 2de). On the test day, mice responded to the test image of the rewarded category A more strongly compared to the test image of the non-rewarded category B (Figure 2ef), but they also responded at a longer latency compared to the train image from category A (Figure 2g). Thus, mice could tell the difference between the two A exemplars. Since licking on the two test images never delivered reward, the mice gradually reduced their licking to both during the course of the session (Figure 2e). To reduce the effect of this within-day learning, we used only the first block on this day, consisting of the first 25% of the trials, for most analyses.

To investigate the patterns of discrimination accuracy across stimuli, we defined a behavioral discrimination index (DI) as the difference in response rates between A and B trials (Figure 2h). On the training trials, the DI was always high with an average of 0.798 ±.019 (mean ± s.e.m.), while on the test trials the average was 0.393 ±.05 (mean ± s.e.m.) (Figure 2i). Thus, similar to our conceptual model (Figure 1), mice nearly always discriminated the training images perfectly, but had varying levels of generalization to the testing images. The DI on test images was nearly always positive (96%) (Figure 2j). For comparison, this level of generalization could only be obtained in an artificial neural network [32] after several layers of computation, indicating that the biological computations may rely on mid-to high-level visual features (Figure 2k).

### Neural generalization

Next, we obtained estimates of neural discrimination accuracy. We did this in a separate set of animals which were presented images passively without task engagement (Figure 3a), thus ensuring that motor variability in behavioral discrimination was uncoupled from the neural responses. While stimuli were presented, we performed large-scale two-photon calcium imaging simultaneously from 10 brain regions and 15,491-73,826 simultaneously-recorded neurons (Figure 3bc, Figure S3). We presented the same eight categories of images as were used in the task, with four different exemplars included from each (Figure 3b). Neurons responded to these images selectively. For example, some neurons responded strongly to individual exemplars, while others responded to multiple exemplars from one category (Figure 3e).

**Figure 3:**
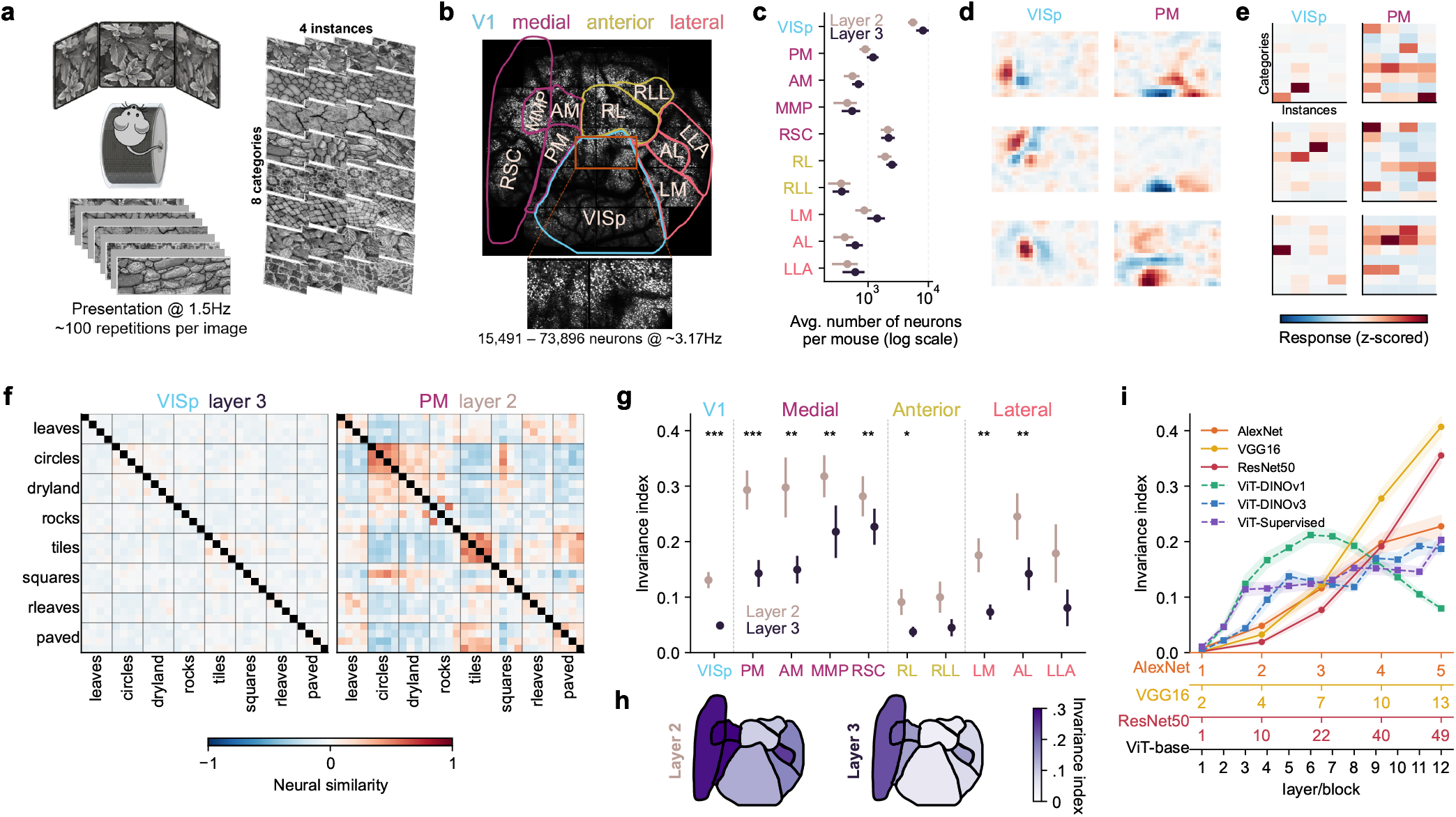
Similarity of neural representations of train and test stimuli. **a**, Schematic of passive stimulus presentation of full-field natural images like those in Figure 2a. (left) and examples of images used (right). **b**, (top) Average two-photon image of recording field of view encompassing 9 primary and higher-order visual areas, and (bottom) a zoom in to illustrate cellular resolution. **c**, Average number of neurons per visual area recorded per mouse and depth. Error bars represent s.e.m across mice (n=12). **d**, Linear receptive fields of three example neurons from VISp and PM **e**, Average neural responses of neurons in **d** to the images in **a. f**, Similarity of vectorized neural representations between all 32 images in **a** for recordings in (left) primary visual cortex at a depth of 250*µ*m and (right) medial visual cortex at a depth of 100*µ*m, averaged across mice (n = 12 mice). **g**, Average invariance index computed between pairs of image categories separately by brain area and depth (n = 12 mice, two-sided paired t-test). Error bars represent s.e.m. across mice. **h**, Summary of invariance indices in **g** visualized on the brain map. **i**, Same as **g** for convolutional neural networks (solid) and the base visual transformer with different loss functions (dashed). Error bars represent s.e.m. across category pairs (n=28).

To obtain a neural discrimination index, we computed the similarity of the neural population vector responses for each recorded region, further divided into layer 2 and layer 3, recorded at 100µm and 250µm from the brain surface respectively (Figure 3f). In some regions, like primary visual cortex (V1), we found an almost uncorrelated neural code across images, while in other regions like posteromedial cortex (PM) we observed similarities that clustered based on the blocks of images coming from the same categories. To determine the level of generalization across images of the same category, we defined an “invariance index” as the average of similarities within-category, minus the similarities across-category (Figure 3gh). Defined this way, invariance was highest in the medial higher-order visual regions, regions that also had the largest receptive field size and lowest spatial frequency preference (Figure S4). Invariance was also higher in layer 2 compared to layer 3, possibly due to long-range horizontal connectivity being more prevalent in layer 2 of cortex [33, 34]. Similar to behavioral generalization, peak neural generalization matched the invariance levels of deeper levels of artificial neural networks (Figure 3i) [32, 35–39].

For comparison, we also recorded responses to the same stimuli in the superior colliculus, and performed the same analyses to find that the invariance index was low, similar to the layer 3 of primary visual cortex (Figure S2).

### Relating behavior to neural activity

Having obtained estimates of both behavioral and neural discrimination to the same set of train and test images, we next related them. For this, we needed to compute a discrimination metric based on the similarity of neural population vectors on the images used in the task (Figure 4a). Similar to the invariance index, the discrimination metric was defined as “within” similarity minus “across” similarity, though in this case the similarities were restricted to just those images used in the behavior (Figure 4bc). On train trials, the “within” similarity was defined as “within” different trial subsets of neural responses to the same image, while on test trials, the “within” similarity was between train and test images of the same image class. Defined this way, the train and test discrimination indices had similar properties to the behavior discrimination matrix, for example, having generally lower values of test DI compared to train DI (Figure 4d). Restricting the analysis to train images, we found little relation between neural and behavioral DI across all brain areas (Figure 4ef), similar to the lack of a relationship in the toy model (Figure 1c). In test trials, we found a relationship between behavior and neural DI as expected from the toy model (Figure 1d) specifically in the medial higher-order visual regions, in both layer 2 and layer 3. This suggests that the medial visual regions are important for the visual transformation from low-level image features to high-level image information that is available for behavior.

**Figure 4:**
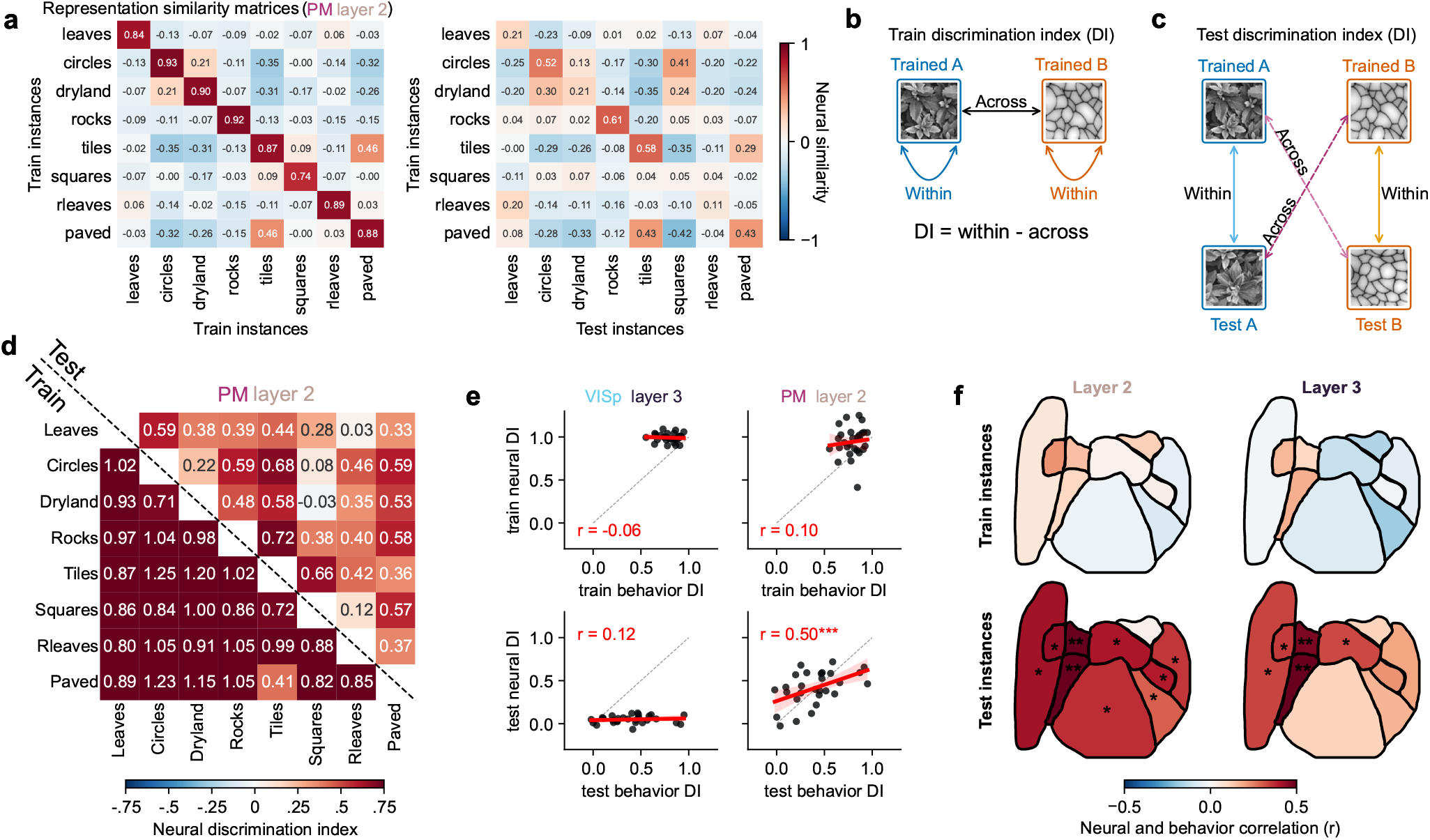
Linking neural and behavioral discrimination. **a**, Similarity of vectorized neural representations between the specific pairs of images used during the task in Figure 2 for layer 2 of VISpm. This includes train/train similarity (left) and train/test similarity (right). **a**, Diagram of the similarities used to compute the train discrimination index in **d. c**, Diagram of the similarities used to compute the test discrimination index in **d. d**, Discrimination indices computed from the similarity matrices in **b** on both train and test trials for layer 2 of VISpm. **e**, Relationship of behavioral and neural discrimination indices (DI) for VISp layer 3 and PM layer 2 on both train (top) and test (bottom) stimuli (Node-based permutation test). **f**, Correlation between behavioral and neural DI on train and test stimuli, divided by brain region and depth (Benjamini-Hochberg corrected p-values from node permutation test).

Finally, we asked whether neural representations in dark-reared mice were also related to behavior in the same way (Figure 5a) [40–42]. First, we verified that neurons in dark-reared mice had similar low-level visual responses as those in normal-reared mice, for example, displaying similar receptive fields (Figure 5b, Figure S5). Then we investigated the responses of neurons in dark-reared mice to the same texture classes (Figure 3a, Figure 5c). Compared to the representations in normal-reared mice, we found a lack of correlation between different stimuli, even in the superficial layers of PM (Figure 5d). The invariance index of these representations was very low across all visual cortical areas (Figure 5e). There was also no significant relation between the neural response patterns and the behavioral response patterns from the normal-reared mice (Figure 5fg). Thus, the relation between neural activity and behavior required visual experience to develop.

**Figure 5:**
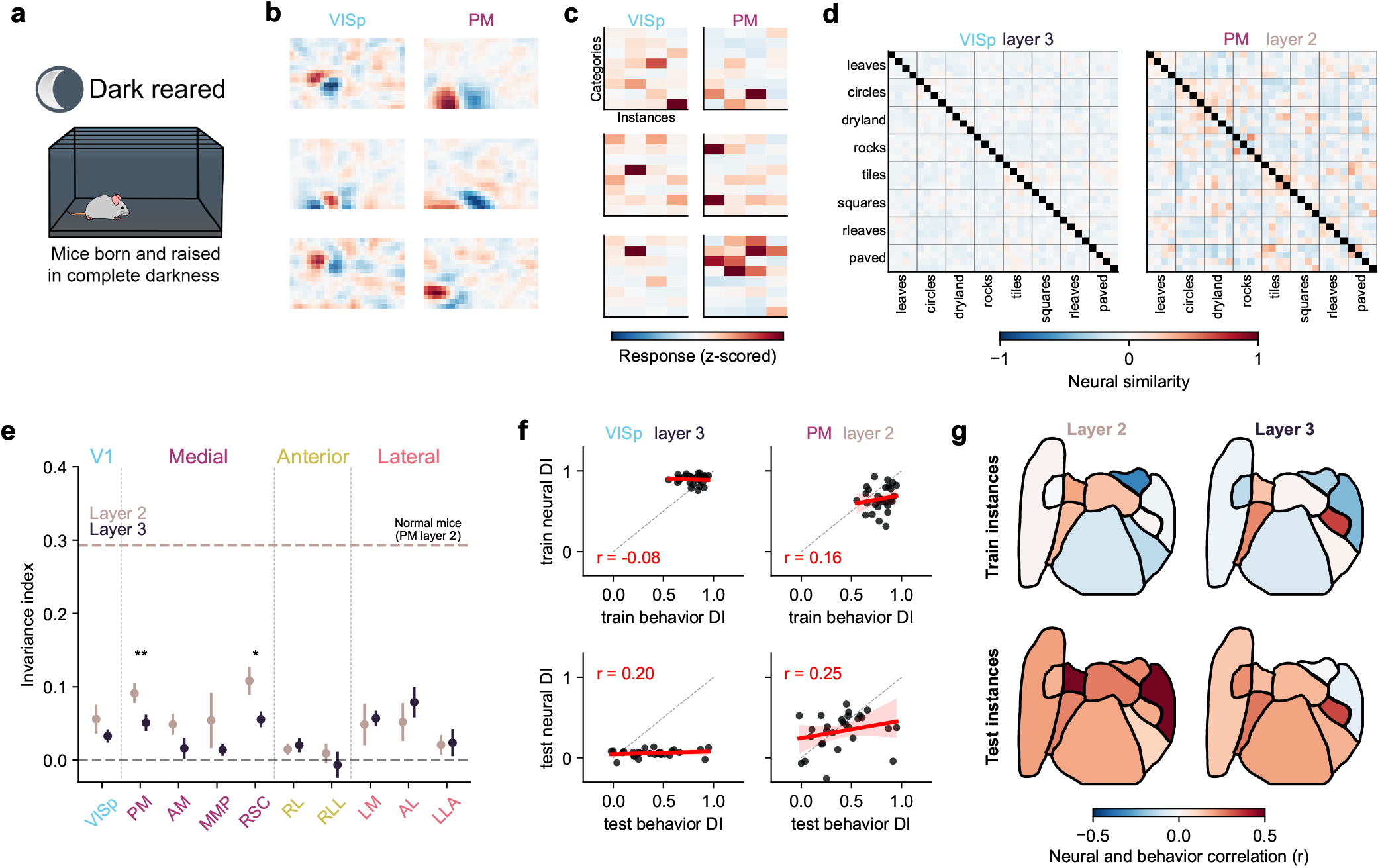
Relation between generalization behavior and neural activity in dark-reared mice. **a**, Illustration of dark-rearing. **b**, Linear receptive fields of three example neurons from VISp and VISpm of dark reared mouse. **c**, Example neural responses of the neurons in **b**, to the images in Figure 3a. **d**, Similarity of vectorized neural representations between all 32 images in Figure 3a for recordings in (left) primary visual cortex at a depth of 250*µ*m and (right) medial visual cortex at a depth of 100*µ*m, averaged across mice (n = 6 mice). **e**, Invariance index in dark-reared mice computed like in Figure 3g (n = 6 mice, two-sided paired t-test). Error bars represent s.e.m. across mice. **f**, Comparison of behavioral and neural DI on train and test stimuli for example regions VISp layer 3 and PM layer 2 (Node-based permutation test). **g**, Correlation between behavioral and neural DI on train and test stimuli, divided by brain region and depth (Benjamini-Hochberg corrected p-values from node permutation test).

## Discussion

Here we found that neural and behavioral discrimination can be related on test trials but not on training trials. Our results point to specific higher-order visual areas in the mouse (the medial region) as the locus of neural computations that increase invariance to low-level features, while also increasing the ability to generalize across exemplars of the same visual categories. This region required visual experience to develop, suggesting a role for un-rewarded visual learning for the development of behavior-related neural representations, as previously suggested in a different context [43]. Future experiments may causally test the role of the medial HVAs in visually-guided behaviors, though this will require cell-targeted perturbations, as the neurons encoding distinct visual categories are intermingled with each other [44, 45].

We discovered that the medial HVA responses were most invariant to textures—previous studies may have not observed these invariance properties due to differing stimulus sets [46–49], or due to not recording from those regions [50]. We also observed higher invariance in more superficial recordings (“Layer 2”, 100µm below pia), which may be partially explained by laminar differences in myelination (see [33] for anatomical characterization in NHPs). It may also be related to differences in synaptic input distributions across depths; for example, more superficial Layer 2/3 neurons receive inputs from a larger spatial area than deeper Layer 2/3 neurons [34, 51, 52].

Our observations in dark-reared mice build upon prior work showing that natural image responses in V1 neurons are altered by dark-rearing [53–55]. In our work, the effect of dark-rearing on invariance was especially pronounced for the “Layer 2” neurons. Another study has also found large differences in responses properties in upper Layer 2/3 neurons in V1 following development in an altered visual context [56]. Other recent work has discovered that dark-rearing specifically alters gene expression in superficial V1 neurons but not deeper neurons [57, 58].

We have contrasted our approach to the more typical parametric variation employed in experiments that attempt to relate neural and behavioral discrimination. While such experiments can induce more mistakes on “harder” trials, these mistakes may not be failures of sensory computations, as the sensory neural representations are strongly discriminative, even on the “hard” trials [28]. Thus, many recent studies have concluded that choice-related neural activity could be represented separately from sensory-related neural activity, dissociating sensory variability from decision-making signals [22, 26, 59, 60]. Multiple brain regions have been identified where neurons do correlate with choice variability, and these brain regions are the subject of intense current scrutiny [14]. Our hope is that the complementary methods we introduced here can similarly point researchers towards brain regions where behavior-related sensory computations occur. While we have demonstrated this approach in the visual system, other sensory modalities are likely amenable to similar investigations.

## Acknowledgments

This research was funded by the Howard Hughes Medical Institute at the Janelia Research Campus and by the Wellcome Trust (220169/Z/20/Z to SS). From the Vivarium, we thank Jim Cox, Crystall Lopez, Amanda Minisi, Anne Kuzspit, Miriam Rose, Gillian Harris, Sarah Lindo, Alexa Gracias, and the research technicians who provided support for animal breeding, husbandry, surgeries, and behavioral training. From JeT, we thank Jon Arnold, Sam Jager and Steven Sawtelle for help with rig maintenance and upgrades. From MBF Bioscience we thank Georg Jaindl and Boris Djiguemde for scanimage support.

## Data availability

The data will be released upon publication.

## Code availability

The code will be released upon publication.

## Ethics declaration

The authors declare no competing interests.

## Methods

All experimental procedures were conducted according to IACUC, and received ethical approval from the IACUC board at HHMI Janelia Research Campus. For the superior colliculus recordings, all procedures were performed in accordance with the UK Animals (Scientific Procedures) Act 1986, under project and personal licenses approved by the Home Office following institutional ethical review.

### Experimental methods

#### Animals

For the normal reared group (n = 12 mice; aged 6–15 months), mice were housed in a reverse light cycle (12h light/12h dark). Six of these mice were bred to express GCaMP8s (TetO-jGCaMP8s × CamK2a-tTa; JAX 037717, 007004), and six expressed GCaMP6s (TetO-GCaMP6s × CamK2a-tTa; JAX 024742, 007004). The dark reared group (n = 6 mice; aged 6–15 months) were also GCaMP6s-expressing mice (JAX 024742, 007004) but they were born, raised and maintained in complete darkness at all times. All maintenance and experimental procedures for these mice were performed using night vision goggles. For transport between the dark holding room and the mesoscope room, animals were shielded with a blackout curtain to ensure their first exposure to light occurred during the experiment. For task-based experiments (n = 15 mice; aged 6–15 months), we utilized C57BL/6 mice, these animals were implanted with a headbar only without craniotomies.

For the superior colliculus recordings, two male mice were used: one Ai14 reporter line (JAX 007908) and one Vgat-IRES-Cre × Ai14 cross (JAX 016962 and 007908). Mice were 8 and 29 weeks old at the time of surgery and were used for data collection until 38 weeks of age. Animals were maintained on a 12-h light/12-h dark cycle.

#### Surgical procedures

For animal surgeries, we followed the protocol described in [61]. Briefly, mice were anesthetized with isoflurane (1.5− 2%) and administered a cocktail of analgesics and anti-inflammatories, including buprenorphine, carprofen, and dexamethasone. A 4 mm craniotomy was centered over the right primary visual cortex (V1; +0.9 mm AP, +2.6 mm ML from lambda). A custom headbar and a 4+5 mm double-glass window were secured using Vetbond and Calibra Universal Resin Cement to allow for chronic optical access. Post-operative care included 48 hours of monitoring, warm LRS fluids (+5% dextrose), and Carprofen (5 mg/kg) delivered subcutaneously.

For the surgical procedures on the superior colliculus, mice were anesthetized with isoflurane and administered subcutaneous injections of meloxicam (5 mg/kg), buprenorphine (0.05 mg/kg), and dexamethasone (4 mg/kg). The scalp was shaved and disinfected, and local analgesia (bupivacaine, 7 mg/kg) was injected subcutaneously prior to the incision. Protective ophthalmic gel was applied to the eyes, and 5% EMLA cream was applied to the ear bars before securing the animal in a stereotaxic frame. Following scalp reflection and removal of connective tissue, a custom headplate was secured to the skull using Superbond C&B. Body temperature was maintained at 37°C throughout the procedure using a feedback-controlled heating pad, and hydration was maintained via subcutaneous injections of Hartmann’s solution (0.01 ml/g/h). Mice were given three days of post-operative recovery with daily Metacam treatment. A 4 mm circular craniotomy was performed (centered at approximately −4.2 mm AP and 0.5 mm ML from Bregma). We injected 230 nl of AAV2/1.Syn.GCaMP6f.WPRE.SV40 [62] at a final concentration of ∼3.5 ×10^12^ GC/ml into the right SC. Injections were performed at a rate of 2.3 nl every 6 s using a Nanoject II (Drummond). To gain optical access to the posterior SC, which is typically obscured by the transverse sinus, a custom-made implant was used to permanently displace the sinus anteriorly. The implant was fixed to the skull using Vetbond and Superbond C&B.

#### Imaging acquisition

For the cortical recordings, two-photon imaging was performed with a 2p-RAM mesoscope [63, 64], the recording and online Z correction were performed with ScanImage [28, 65], at 3 Hz, and two depths 100 µm (layer 2) and 250 µm (layer 3).

For the superior colliculus recordings, two-photon imaging was performed using a standard resonant microscope (HyperScope, Scientifica) equipped with a 16x, 0.8 NA water immersion objective (N16XLWD-PF, Nikon) and controlled by ScanImage (basic version 2022). Excitation light at 920 nm was delivered by a femtosecond laser (Chameleon Vision II, Coherent). A single plane was imaged at depths of 25–115 µm from the surface of the superior colliculus. The field of view spanned 270–730 µm in both directions at a resolution of 512 x 512 pixels. The acquisition frame rate was 30 Hz.

During the recordings, mice were head-fixed and positioned centrally atop a metallic, plastic, or styrofoam cylindrical treadmill to allow for voluntary locomotion [66]. A rotary encoder was used to record running speed.

All animals were handled using refined techniques for at least three days prior to head-fixation acclimation. We implemented a standardized pre-training pipeline: mice were handled for 30 minutes daily and gradually acclimated to the recording rig and head fixation in 20-minute increments until they exhibited no signs of distress.

#### Visual stimuli and experimental design

For the cortical recordings, stimuli were presented using three LED displays (4:3 aspect ratio) arranged to provide a surrounding field of view covering 270° in azimuth. To prevent direct light contamination of the PMTs, green-light-excluding filters were placed in front of the screens, and stimulus rendering was restricted to the blue and red channels. Stimuli were presented using Psychtoolbox in Matlab [67].

For the superior colliculus recordings, stimuli were presented on a single monitor covering −149° to −57° azimuth and −32° to +48° elevation, with brightness levels linearized around a mean luminance of 13 cd/m^2^. Stimuli was presented using BonVision [68] to correct for geometric distortion and maintain uniform stimulus scaling.

We created a dataset of grayscale images consisting of eight texture categories with four unique images per category (Figure 3a). These images were of size 66×264 pixels, and thus each pixel in the image was ∼1 degree of visual angle. The four unique exemplars were generated by randomly cropping eight different high-resolution texture images at random locations, orientations, and spatial scales. To isolate invariant coding from low-level features, we matched the Fourier spectra across the eight categories. To do this, we calculated the global average amplitude spectrum across all images and normalized the mean amplitude spectrum of each category to this reference.

In passive recordings, we presented the visual stimuli to the mice interleaved with gray screen, at a rate of ∼ 1.5 Hz (Figure 3a). In one recording session, the 32 texture images were presented several times in randomized order each time: 82±22 trials for the normal-reared mice, 55±28 trials for the dark-reared mice, and 100 trials for the SC mice. In half of the normal-reared mice and in all of the dark-reared mice we also presented up to 15,000 different natural images in grayscale [69], interleaved with the 32 images, during this recording session, which we used for retinotopic mapping and for fitting Gabor models to each neuron. In the other six normal-reared mice, we recorded a separate imaging session in which we presented 500 different natural images repeated three times to use for retinotopic mapping.

In the task-experiments, virtual pseudo-corridors were constructed using images from a 32-texture set. Each corridor spanned 400 cm in total, consisting of a 300 cm stimulus zone followed by a 100 cm gray-screen inter-trial interval. Stimulus visibility was controlled by linearly modulating the contrast via the alpha channel: alpha increased over the first 150 cm and decreased over the subsequent 150 cm (Δα = 0.0066 per cm). If locomotion exceeded a threshold of 6 cm/s, then the virtual pseudo-corridor moved forward at a constant speed of 57.1 cm/s, regardless of the absolute speed of the mouse. This configuration normalized the traversal of the 400 cm corridor to ∼ 7 seconds when the mouse runs continuously.

#### Online Gaze Correction

In the passive stimulus presentation recordings, we performed real-time tracking of the left pupil using a customized version of Facemap [70]. To maintain consistent retinal input, horizontal eye position shifts triggered a synchronous adjustment of the stimulus center-point, ensuring stimuli remained centered on the horizontal axis of rotation of the eye.

#### Water restriction procedure

Water restriction was conducted in accordance with IACUC guidelines. Restricted animals received a target of 1.0 ml of water per day, with adjustments ranging from 0.7 to 1.5 mL based on individual performance and health status. Mice were gradually introduced to the restriction goal by reducing their intake by 0.5 mL daily over the three days preceding the start of experiments (from 2.0 mL down to 1.0 mL). During the task, mice could obtain up to 0.7 mL of water if all 200 available rewards were earned (3.5 µL per reward). Any remaining volume (total daily allowance minus the amount received during the experiment) was provided one hour after the training session. Throughout the restriction period, body weight, physical appearance, and behavior were monitored daily using a standard quantitative health assessment system [71].

#### Water reward delivery and lick detection

A capacitance detector was connected to a metal lick port to monitor licking behavior. During training sessions, the mouse received a water reward (3.5 µL) if it licked anywhere in the designated reward region for the appropriate stimulus category (“train A”), and the reward was delivered immediately after the first detected lick in the reward region. In test sessions, if the mouse did not lick on a “train A” trial, then a “passive reward” was delivered at the start of the next inter-trial interval to maintain animal engagement, these instances were recorded as performance errors.

#### Behavioral training

After head-fixation acclimatization, mice underwent a five-day running regimen (1 hour/day) to ensure smooth and continuous locomotion on the cylindrical treadmill. During this phase, water rewards were triggered by maintaining a target running speed for a specified duration; these thresholds were adjusted *ad hoc* for each mouse based on performance.

Mice were divided into two experimental cohorts (Figure S1):

- Pilot group: Animals were trained until they achieved three days of a discrimination index (DI) greater than 0.7, which ranged from 5 - 15 days of training, and then tested with the new images.
- Homogeneous training group: Animals trained for exactly four days and tested on the fifth (this subset of sessions were used for Figure 2e, all other analyses used both groups).

## Data analysis

For analysis, we used Python 3 [72], primarily based on numpy, scipy, statsmodels, scikit-learn and pytorch [73–77]. Figures were made using matplotlib, seaborn and jupyter-notebook [78–80].

### Behavioral discrimination index

To quantify categorical discrimination, we calculated a behavioral Discrimination Index (*DI*) for both training and test sessions (Figure 2hi). For training sessions, the index (*DI*_*train*_) was defined as:

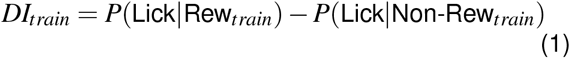

where *P*(Lick| Rew) / *P*(Lick| Non-Rew) are the proportions of rewarded / non-rewarded trials with at least one lick in the second half of the corridor, where licks trigger rewards on rewarded trials. The performance during test trials was similarly calculated and normalized by the training performance of the same session to yield the Test DI (*DI*_*test*_):

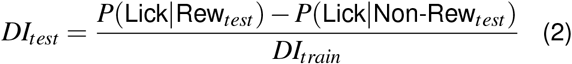

This normalization procedure accounts for session-to-session variability in motivation and ensures that test performance is evaluated relative to the animal’s baseline discrimination proficiency.

Test session analyses were restricted to the first block of trials (25% of the total), to reduce the effect of within-day learning.

#### Processing of calcium imaging data

Calcium imaging data were processed using the Suite2p pipeline (v0.9.3) [81] (available at www.github.com/MouseLand/suite2p). As previously described [21], Suite2p was utilized for rigid and non-rigid motion correction, automated region of interest (ROI) detection, cell classification, and neuropil subtraction. Spike deconvolution was performed using a non-negative algorithm with a fixed decay timescale of 0.75 s [82, 83]. All analyses were conducted using the resulting deconvolved fluorescence traces. The response latency between stimulus presentations and neural responses was determined by the time lag which maximized the population signal variance (*c*_*sig*_, see below). On average across mice, this latency was one frame, and so neural responses were defined as the activity on the first frame after each stimulus presentation.

#### Single-neuron response reliability

We restricted our analyses to the top 10% most stimulus-driven neurons within each ROI (defined by area and layer). Reliability was quantified using the signal variance, estimated by an approach based on [84] calculating the trial-to-trial covariance across repeated presentations of the identical stimuli from the 32 texture image set:

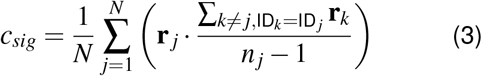

where **r**_*j*_ is the response vector to trial *j, n*_*j*_ is the total number of repeats for that stimulus, and *N* is the total number of trials. **r**_*j*_ · 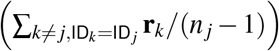, represents the product of the response in trial *j* and the mean response of all other trials sharing the same stimulus identity.

By using this leave-one-out cross-covariance approach, we ensure that the resulting signal variance estimate (*c*_*sig*_) is not inflated by trial-specific noise, as independent noise components across trials *j* and *k* have an expected product of zero. Consequently, *c*_*sig*_ specifically captures the portion of the neural variance that is reliably evoked by the stimulus. This approach isolates the consistent stimulus-evoked component from independent trial-to-trial noise.

Prior to analysis, responses for each neuron were *z*-scored across all stimulus presentations from the 32 texture image set, to normalize for differences in firing rates across the population.

#### Invariance index

To quantify the category selectivity of neural populations, we calculated an invariance index for each area and depth per mouse, for each pair of texture categories (Figure 3g). First we computed the trial-averaged responses of the neurons to the 32 texture image set, denoted as vectors across neurons **r**, from different categories, i.e. *a* and *b*, and different exemplars *i* and *j*. Then we used these population response vectors to compute the invariance index (*I*_*index*_):

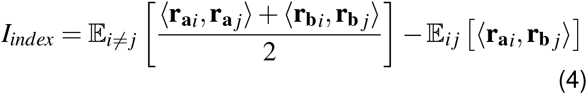

where, ⟨·, ·⟩ is defined as the Pearson correlation between population response vectors. The first term represents the mean similarity between distinct instances (*i ≠ j*) of the same category (within-category similarity), while the second term represents the mean similarity between instances of different categories (between-category similarity). A positive *I*_*index*_ indicates that neural representations are more stable across transformations of the same category than they are across different categories, reflecting the emergence of category-invariant representations within the visual hierarchy. This is computed for all category pairs and averaged.

Due to the relatively low neuron counts in Superior Colliculus recordings, we concatenated the top 10% most reliable units from the three sessions. The invariance index and its associated variability were then estimated via bootstrapping the concatenated population (10,000 iterations).

#### Neural discrimination index

To measure the discriminability of neural representations for the specific stimuli used during the task, we calculated a neural discrimination index (Figure 4b-d). For training stimuli, we employed a split-half correlation approach to account for trial-to-trial variability. The training index 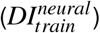 was defined as:

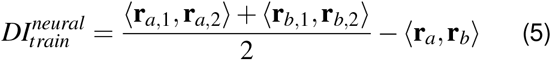

where ⟨**r**_*x*,1_, **r**_*x*,2_⟩ represents the Pearson correlation between the mean population response vectors of two independent halves of the trials for stimulus *x*. The term ⟨**r**_*a*_, **r**_*b*_⟩ denotes the correlation between the mean responses to the training stimuli of category *a* and *b*.

To evaluate the representation of novel stimuli relative to the learned categories, the test discrimination index 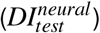 was calculated as:

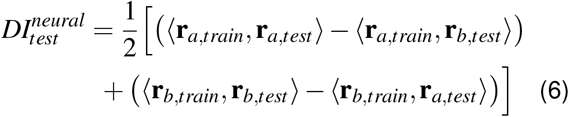

where⟨**r**_*x,train*_, **r**_*y,test*_⟩ is the correlation between the training stimulus of category *x* and the test stimulus of category *y*. Finally, the 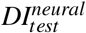 was normalized by the 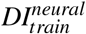 of the given category pair, in a similar fashion to its behavior counterpart to make possible the direct comparison.

#### Retinotopy

Spatial receptive locations were estimated following the procedure described by [43], using the passive recording sessions described above. Initially, spatial kernels (*n* = 200) were fitted to the neural responses of a reference mouse [85]. This model was subsequently applied to recordings from all other subjects to identify the preferred kernel and corresponding spatial location for each cell. Finally, these locations were aligned to the reference mouse’s spatial map, where the recorded regions had been segmented based on a computed sign map [86]. Results from this retinotopic mapping are plotted in Figure S3 and Figure S4 for normal-reared and dark-reared mice, respectively.

#### Linear receptive fields

Linear receptive fields (Figure 3d, Figure 5b) were estimated using linear regression, fit to predict the z-scored neuron responses from the visual stimuli downsampled by a factor of 4, with a regularization constant found via optimization of the mean squared error.

#### Artificial neural network analysis

To compare the neural invariance index with invariance in artificial vision models, we applied the same metric to layer-wise representations from six pretrained networks (Figure 3i). Specifically, we analyzed AlexNet [32], VGG16 [35], and ResNet50 [36], all trained in a supervised manner on ImageNet [87], as well as three Vision Transformer (ViT) models, all of the ViT-Base/16 variant (12 transformer blocks, hidden dimension 768, 16 × 16 patches): a supervised ViT-B/16 [37], DINOv1, and DINOv3 [39]. All models were loaded with frozen pretrained weights and run in evaluation mode.

To match the aspect ratio of our texture stimuli while ensuring compatibility with the ViT patch size, all input images were resized to 64× 256 pixels (divisible by the patch size of 16, yielding 4 ×16 = 64 patch tokens per ViT image). For the CNNs, images were normalized using the standard ImageNet statistics (mean = (0.485, 0.456, 0.406), std = (0.229, 0.224, 0.225)). For the ViTs, images were normalized using each model’s default HuggingFace image processor (ImageNet statistics for DINOv1 and DINOv3; mean = std = (0.5, 0.5, 0.5) for the supervised ViT).

For each CNN, we extracted representations at five architectural blocks rather than at every individual convolution. In AlexNet, each block corresponded to one of the five convolutional layers (C1–C5), taken after ReLU (and after max-pooling for C1, C2, and C5). In VGG16, the 13 convolutions were grouped into five max-pool stages containing 2, 2, 3, 3, and 3 convolutions, respectively. In ResNet50, the five blocks were comprised of the initial stem convolution and the four subsequent residual stages. For the ViTs, we extracted the activations of the patch tokens after each of the 12 transformer blocks. For both the CNN and ViT models, these per-block activations were treated as the population-response analogue of **r**, and the invariance index *I*_*index*_ was computed independently at each block.

#### Nearest-Neighbor layer-wise decoding analysis

We evaluated the categorical information preserved in AlexNet at each stage of the model hierarchy using a 1-nearest neighbor (1-NN) decoding procedure on pairs of categories (Figure 2k) (chance is 50%). We used the same train instances and test instances used in the behavior task. As described above, images were normalized using the standard ImageNet statistics, and the activations were extracted after each Alexnet layer. We then computed the pairwise Cosine distance between the activations of the two training images and the test images. The 1-NN decoder assigned each test image to the category of the nearest training image. The decoding analysis was performed independently for each convolutional layer (Cnv1–Cnv5) and the fully connected layer (FC).

#### Gabor model

The Gabor model was fit to neural responses to natural images from six recording sessions in six of the normal-reared mice. In these sessions there were 3,439–9,380 unique training images presented, and 410–500 test images with two repetitions per image presented. Images were downsampled by a factor of two for model fitting. The repeated test presentations were used to estimate explainable variance and the fraction of explainable variance explained (FEVE) [89]. We used the same model as in [84]. In brief, we constructed 7,776 Gabor filters with the following parameters: spatial frequency in cycles per degree (0.05, 0.10, 0.15, 0.20, 0.25, 0.30, 0.35, 0.40, 0.45), orientation (0, π/8, π/4, 3^*^π/8), phase (0, π/2, π, 3^*^π/2), standard deviation σ in degrees (2, 4, 6, 8, 10, 12, 14, 16, 18), and eccentricity β (1, 1.5, 2). These filters were applied convolutionally to all the images, and then the simple and complex cell responses (using phase-offset filters) were computed for each filter and location, with a ReLU nonlinearity. We fit each neuron as a linear combination of the simple and complex cell responses. For each neuron, the parameter set and location that maximized the variance explained on the training images was chosen as the optimal Gabor filter. This simple/complex Gabor model achieved a mean FEVE of 0.175 on the top 10% signal variance neurons pooled across all visual areas (n = 31,401 neurons from 6 mice). The receptive field size was defined as the area of the ellipse of the Gabor filter at one standard deviation 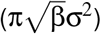.

#### Statistics and reproducibility

We performed paired, two-sided t-tests in figure panels 2f, 3g, 5g, and S4cd; one sample two-sided t-test in 2i; and a Cramér-von Mises test in 2g. No adjustments were made for multiple comparisons on these tests.

To quantify the relationship between neural representations and behavior, we calculated the Pearson correlation (*r*) between the neural discrimination matrix and the behavioral discrimination matrix for each ROI (defined by area and layer). Because these matrices were constructed from all 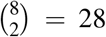 pairwise combinations of the eight stimulus categories, the resulting data points were non-independent. To account for this dependency and preserve the intrinsic structure of the stimulus space, we employed a node-based permutation approach. For each of 10,000 iterations, we randomly shuffled the category labels of the behavioral discrimination matrix, thereby preserving its underlying structure and first-order statistics while breaking the specific link to neural activity. Non-parametric *p*-values were determined as the proportion of permutations where the null correlation coefficient (*r*_*null*_) was greater than or equal to the observed coefficient (*r*_*obs*_). To control for the false discovery rate across the 20 recorded ROIs, raw *p*-values were adjusted with the Benjamini-Hochberg (BH) procedure, with a significance threshold of *p <* 0.05. The observed coefficient and the raw *p*-values are shown for two example ROIs in Figure 4e and Figure 5f while the BH corrected *p*-values are shown in cortical maps in Figure 4f and Figure 5g.

For all figures * denotes *p <* 0.05, ** denotes *p <* 0.01, and *** denotes *p <* 0.001, and error bars represent the standard error of the mean, the exact p-values for each test are listed below:

- 2f train: 7.34 × 10^−46^, test: 2.79 × 10^−11^
- 2g: 6.79 × 10^−10^
- 2i: 9.43 × 10^−9^
- 3g:
  – VISp: 8.86× 10^−6^
  – VISpm: 4.68 × 10^−5^
  – VISam: 0.001
  – VISmmp: 0.005
  – RSC: 0.002
  – VISrl: 0.012
  – VISrll: 0.141
  – VISlm: 0.003
  – VISal: 0.004
  – VISlla: 0.133
- 4f train instances:
  – layer 2 (raw / corrected):
    * VISp: 0.552 / 0.760
    * VISpm: 0.297 / 0.760
    * VISam: 0.225 / 0.760
    * VISmmp: 0.125 / 0.760
    * RSC: 0.358 / 0.760
    * VISrl: 0.461 / 0.760
    * VISrll: 0.26 / 0.760
    * VISlm: 0.646 / 0.760
    * VISal: 0.418 / 0.760
    * VISlla: 0.605 / 0.760
  – layer 3 (raw / corrected):
    * VISp: 0.627 / 0.760
    * VISpm: 0.185 / 0.760
    * VISam: 0.301 / 0.760
    * VISmmp: 0.197 / 0.760
    * RSC: 0.510 / 0.760
    * VISrl: 0.725 / 0.805
    * VISrll: 0.777 / 0.817
    * VISlm: 0.817 / 0.817
    * VISal: 0.522 / 0.760
    * VISlla: 0.601 / 0.760
- 4f test instances:
  – layer 2 (raw / corrected):
    * VISp: 1.08 × 10^−3^ / 0.021
    * VISpm: 4 × 10^−4^ / 0.002
    * VISam: 4 × 10^−4^ / 0.002
    * VISmmp: 3.8 × 10^−3^ / 0.012
    * RSC: 4.4 × 10^−3^ / 0.012
    * VISrl: 4.5 × 10^−3^ / 0.012
    * VISrll: 0.442 / 0.442
    * VISlm: 0.031 / 0.045
    * VISal: 7 × 10^−3^ / 0.017
    * VISlla: 8.8 × 10^−3^ / 0.019
  – layer 3 (raw / corrected):
    * VISp: 0.257 / 0.270
    * VISpm: 4 × 10^−4^ / 0.002
    * VISam: 4 × 10^−4^ / 0.002
    * VISmmp: 0.023 / 0.038
    * RSC: 0.0183 / 0.033
    * VISrl: 0.03 / 0.045
    * VISrll: 0.223 / 0.248
    * VISlm: 0.176 / 0.207
    * VISal: 0.084 / 0.105
    * VISlla: 0.084 / 0.105
- 5e:
  – VISp: 0.100
  – VISpm: 0.006
  – VISam: 0.064
  – VISmmp: 0.289
  – RSC: 0.023
  – VISrl: 0.335
  – VISrll: 0.541
  – VISlm: 0.755
  – VISal: 0.450
  – VISlla: 0.831
- 5g train instances:
  – layer 2 (raw / corrected):
    * VISp: 0.683 / 0.878
    * VISpm: 0.203 / 0.752
    * VISam: 0.150 / 0.752
    * VISmmp: 0.499 / 0.878
    * RSC: 0.491 / 0.878
    * VISrl: 0.222 / 0.752
    * VISrll: 0.969 / 0.969
    * VISlm: 0.749 / 0.878
    * VISal: 0.488 / 0.878
    * VISlla: 0.551 / 0.878
  – layer 3 (raw / corrected):
    * VISp: 0.659 / 0.878
    * VISpm: 0.107 / 0.752
    * VISam: 0.225 / 0.752
    * VISmmp: 0.743 / 0.878
    * RSC: 0.574 / 0.878
    * VISrl: 0.470 / 0.878
    * VISrll: 0.790 / 0.878
    * VISlm: 0.444 / 0.878
    * VISal: 0.032 / 0.648
    * VISlla: 0.869 / 0.915
- 5g test instances:
  – layer 2 (raw / corrected):
    * VISp: 0.115 / 0.195
    * VISpm: 0.068 / 0.195
    * VISam: 0.007 / 0.076
    * VISmmp: 0.146 / 0.195
    * RSC: 0.124 / 0.195
    * VISrl: 0.083 / 0.195
    * VISrll: 0.074 / 0.195
    * VISlm: 0.223 / 0.252
    * VISal: 0.109 / 0.195
    * VISlla: 0.005 / 0.076
  – layer 3 (raw / corrected):
    * VISp: 0.108 / 0.195
    * VISpm: 0.045 / 0.195
    * VISam: 0.211 / 0.252
    * VISmmp: 0.069 / 0.195
    * RSC: 0.226 / 0.252
    * VISrl: 0.146 / 0.195
    * VISrll: 0.481 / 0.506
    * VISlm: 0.131 / 0.195
    * VISal: 0.032 / 0.195
    * VISlla: 0.617 / 0.617

---

**S1:**
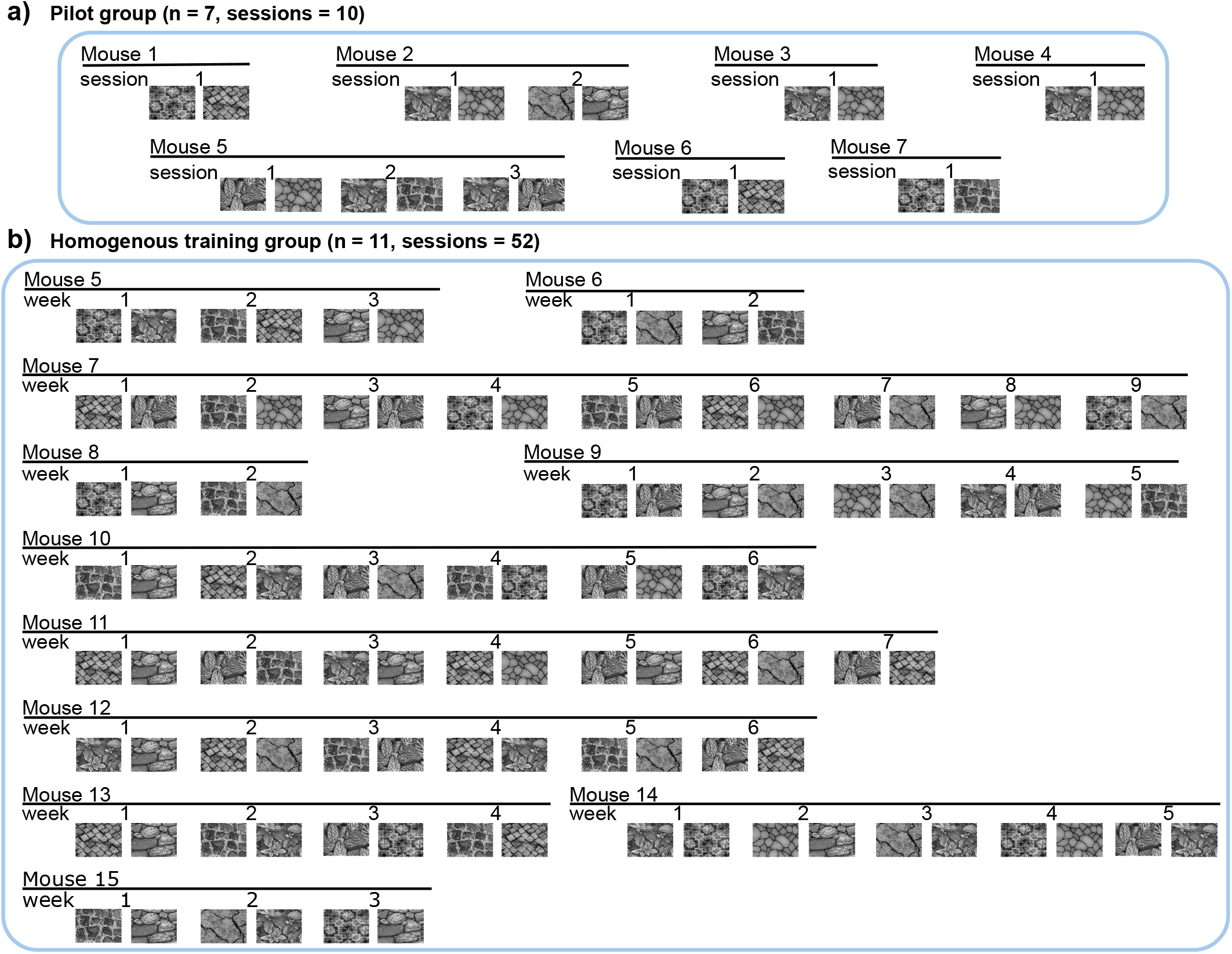
Task experimental cohorts. **a** Pilot group. **b** Homogeneous training group

**S2:**
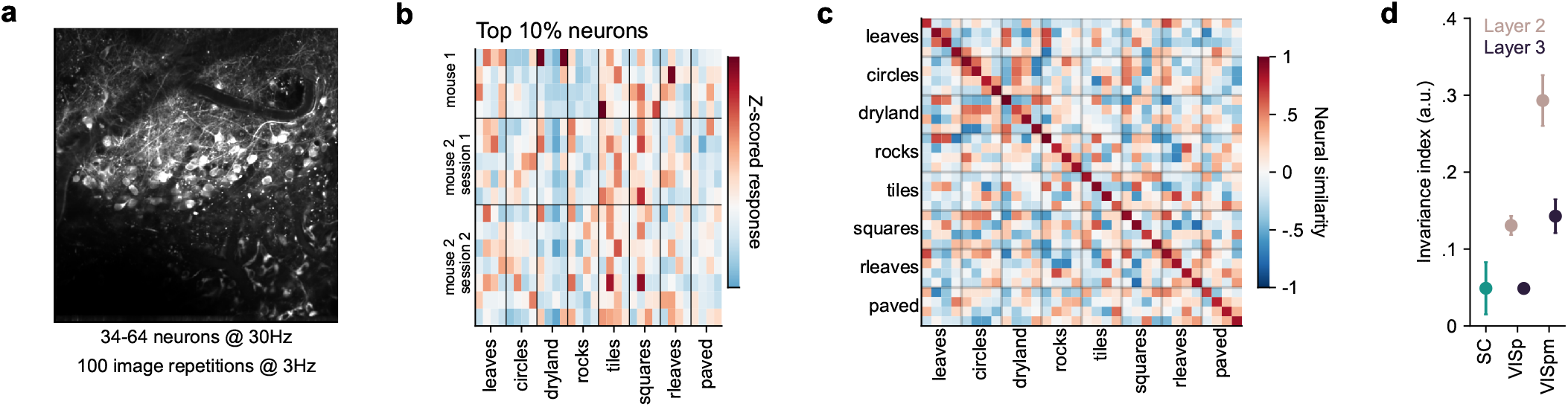
Responses in superior colliculus (SC) to texture images. **a**, Mean image from example recording. **b**, Trial-averaged responses of the top 10% most reliable neurons across three sessions in two mice. **c**, Similarity between responses to texture images (as in Figure 3f), using neurons in **b. d**, Comparison of invariance indices between SC and visual cortical areas (from Figure 3g).

**S3:**
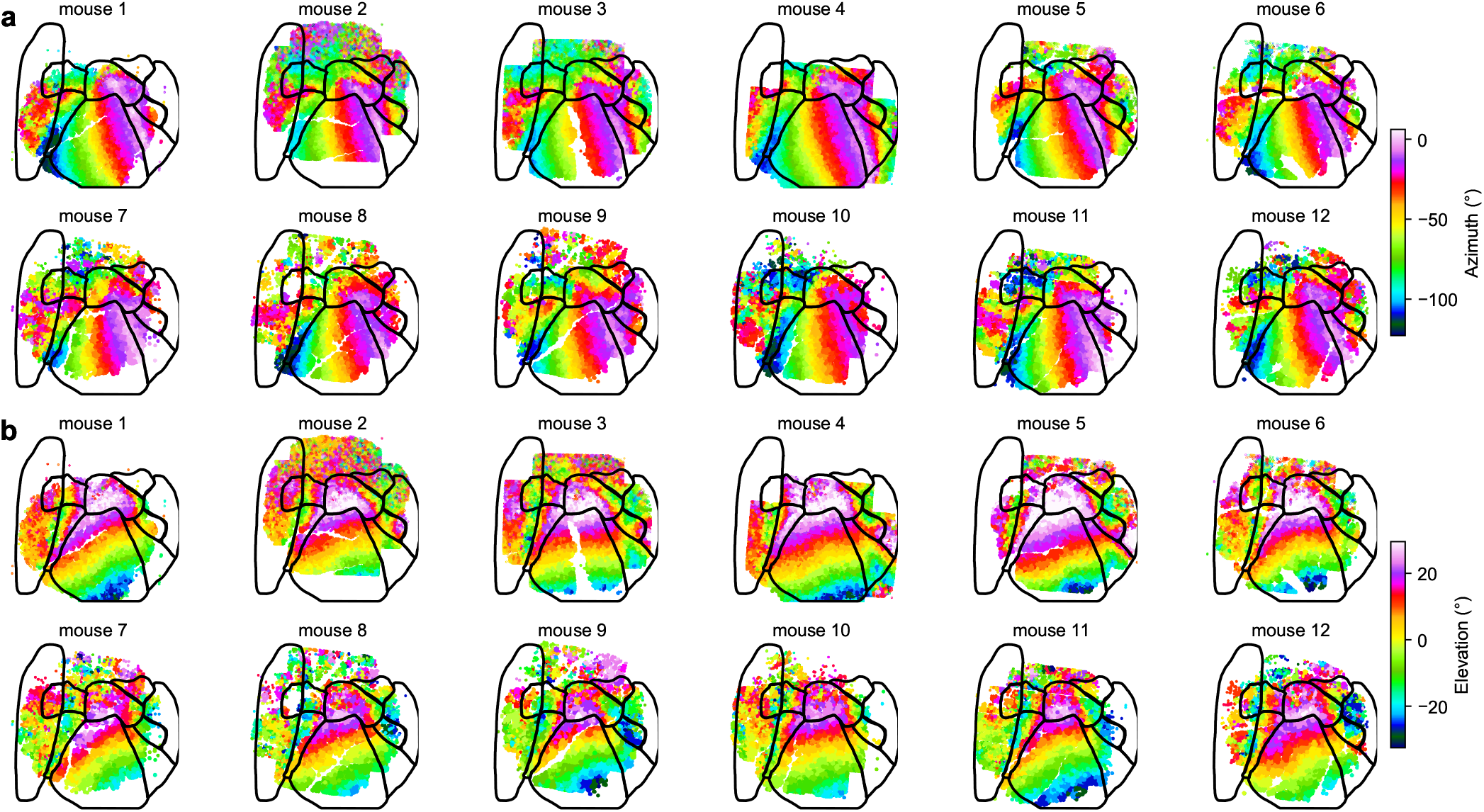
Retinotopic maps in the normal reared group. **a-b**, Each neuron recorded is colored by its preferred visual angle in azimuth (a) and elevation (b).

**S4:**
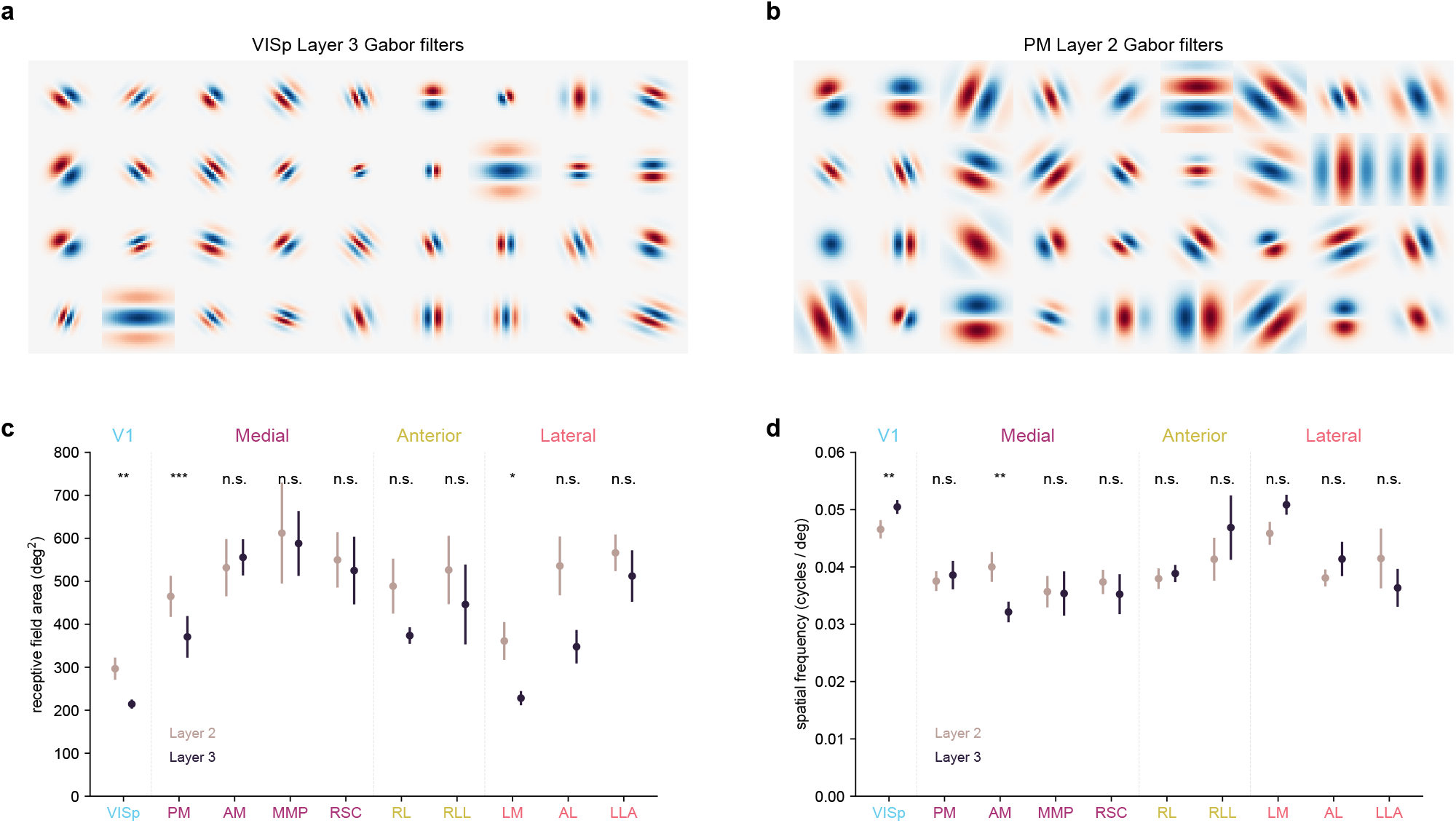
Gabor model fit to cortical neurons. **a**, Example Gabor filters fit to VISp neurons in Layer 3. **b**, Same as **a** for PM neurons in Layer 2. **c**, Quantification of receptive field area from Gabor fits across areas and layers. Error bars represent s.e.m. across mice (n=6). **d**, Same as **c** for spatial frequency.

**S5:**
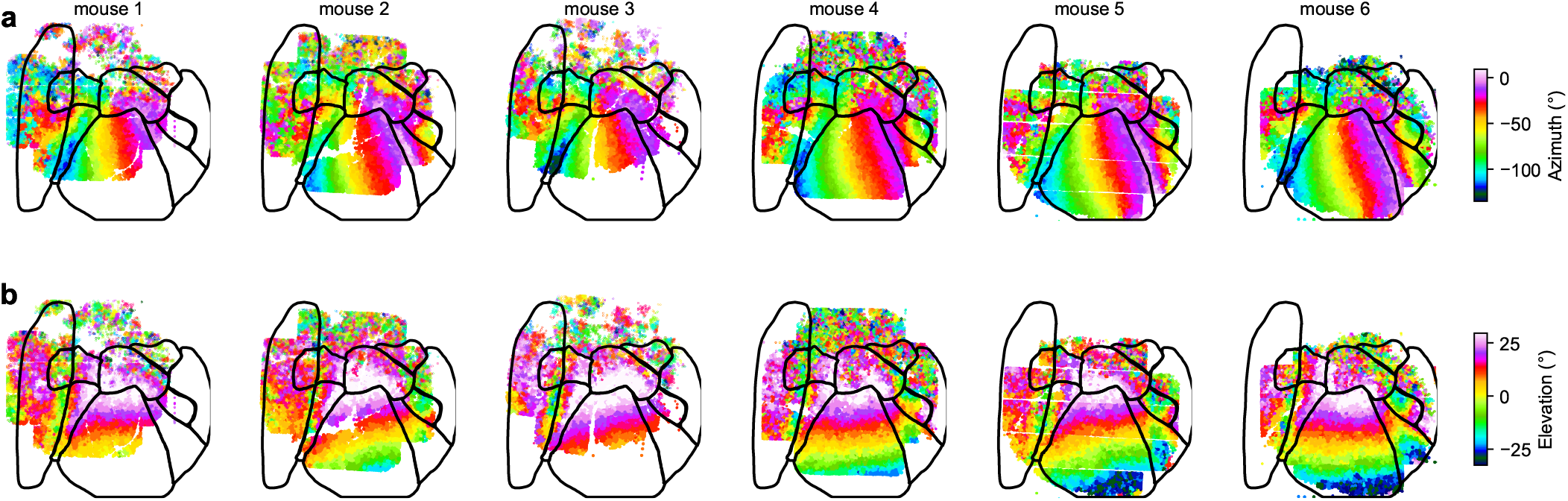
Retinotopic maps in the dark reared group. **a-b**, Each neuron recorded is colored by its preferred visual angle in azimuth (a) and elevation (b).

